# Analysis of distance-based protein structure prediction by deep learning in CASP13

**DOI:** 10.1101/624460

**Authors:** Jinbo Xu, Sheng Wang

## Abstract

This paper reports the CASP13 results of distance-based contact prediction, threading and folding methods implemented in three RaptorX servers, which are built upon the powerful deep convolutional residual neural network (ResNet) method initiated by us for contact prediction in CASP12. On the 32 CASP13 FM (free-modeling) targets with a median MSA (multiple sequence alignment) depth of 36, RaptorX yielded the best contact prediction among 46 groups and almost the best 3D structure modeling among all server groups without time-consuming conformation sampling. In particular, RaptorX achieved top L/5, L/2 and L long-range contact precision of 70%, 58% and 45%, respectively, and predicted correct folds (TMscore>0.5) for 18 of 32 targets. Although on average underperforming AlphaFold in 3D modeling, RaptorX predicted correct folds for all FM targets with >300 residues (T0950-D1, T0969-D1 and T1000-D2) and generated the best 3D models for T0950-D1 and T0969-D1 among all groups. This CASP13 test confirms our previous findings: (1) predicted distance is more useful than contacts for both template-based and free modeling; and (2) structure modeling may be improved by integrating alignment and co-evolutionary information via deep learning. This paper will discuss progress we have made since CASP12, the strength and weakness of our methods, and why deep learning performed much better in CASP13.

## Introduction

Significant progress has been achieved on protein structure prediction due to the development of two major ideas: (1) direct coupling analysis (DCA) for co-evolution analysis^1-5^; and (2) very deep and fully convolutional residual neural network (ResNet) for protein contact and distance prediction^6, 7^. DCA may recover a small set of long-range native contacts when the protein under study has a large number of sequence homologs. In contrast, deep ResNet not only works very well on proteins without many sequence homologs, but also can directly predict inter-residue or inter-atom distance.

In CASP12 and previous CAMEO tests we have demonstrated that deep ResNet can greatly improve contact prediction^6, 8-10^ and that even without time-consuming conformation sampling, contacts predicted by deep ResNet can result in correct folding of (even membrane) proteins without detectable homology in PDB^11^. Afterwards, the power of deep convolutional neural network has been further validated by other research groups who have reimplemented similar deep networks for contact prediction^12-14^. Although contact prediction itself is an important problem that needs further research, we have switched our focus from contact to distance prediction and accordingly distance-based protein structure modeling. This is because a distance matrix contains finer-grained information than a contact matrix and provides more physical constraints of a protein structure, e.g., distance is metric while contact is not. That is, a distance matrix can determine a protein structure much more accurately than a contact matrix. Trained by distance instead of contact matrices, ResNet may automatically learn more about the intrinsic properties of a protein structure and thus, greatly reduce the conformation space, improve folding accuracy and shorten running time needed for protein folding.

Although not totally new, there were only few studies on protein distance prediction and their accuracy were not very satisfactory^15-18^. In 2012, we have employed a probabilistic neural network to predict inter-residue distance distribution from sequence profile and mutual information and then from predicted distribution derived protein-specific distance-based potential for decoy ranking^19^, remote homology detection^20^ and folding simulation^21^. In these studies, we have shown that protein-specific distance potential derived from machine learning performs well on decoy ranking and remote homology detection and comparably with other methods in folding simulation.

Prior to CASP13, we have extended our deep ResNet to protein distance prediction and showed that distance-based potential predicted by ResNet may significantly improve protein threading for targets without good templates in PDB^22^. We have also implemented a simple and efficient distance geometry algorithm that may quickly fold a protein sequence from distance and torsion angles predicted by deep ResNet^23^. Our deep ResNet not only can predict distance matrix from sequence and co-evolutionary information, but also from template and alignment information. In this paper we describe our methods for distance prediction and distance-based protein threading and folding, analyze our performance in CASP13 and discuss the strengths and weaknesses of our approach. We will also examine a few specific targets and highlight our views on the future development and challenge.

## Materials and Methods

### Deep dilated ResNet for protein distance and contact prediction

We use a very similar deep ResNet as described in our previous paper^6^ to predict the Euclidean distance distribution of two atoms (of different residues) in a protein to be folded. Our ResNet model consists of one 1D deep ResNet, one 2D deep dilated ResNet and one Softmax layer (Fig. 1). The 1D and 2D ResNets capture long-range sequential and pairwise context, respectively. The 2D ResNet used a dilated instead of the traditional convolutional operation^24^ to yield slightly better accuracy with fewer model parameters. The 1D and 2D ResNets use ∼7 and ∼60 convolutional layers, respectively, and kernel size of 15 and 5×5, respectively.

**Figure 1.**
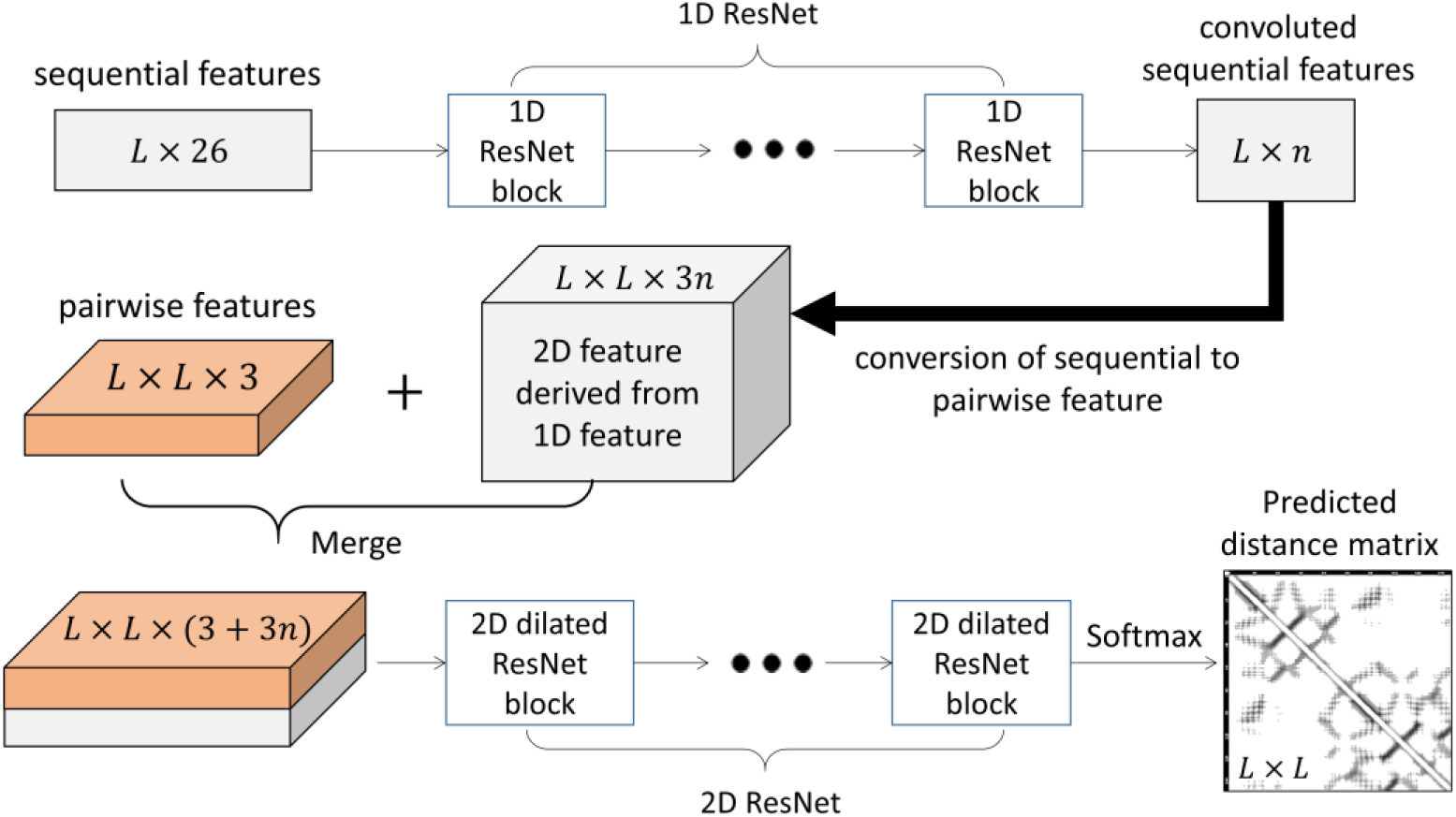
The overall architecture of deep dilated ResNet used in CASP13.

We discretize inter-atom distance into 25 bins: <4.5Å, 4.5-5Å, 5-5.5Å, …, 15-15.5Å, 15.5-16Å, and >16Å and treat each bin as a label for classification. The ResNet model for distance prediction is trained using the same procedure as before^6^. Contact prediction is fulfilled by summing up the probability of all the C_β_-C_β_ distance bins falling into interval [0, 8Å]. Our distance-based contact prediction has 3-4% better long-range prediction precision than the ResNet model directly trained from contact matrices. Besides C_β_-C_β_ distance distribution, we also trained individual ResNet models to predict distance distribution for the following atom pairs: C_α_-C_α_, C_α_-C_g_, C_g_-C_g_, and N-O. Here C_g_ represents the first CG atom in an amino acid. When CG does not exist, OG or SG is used. The predicted distance of these 5 atom pairs is used together to fold a protein, which on average is slightly better than using the predicted C_β_-C_β_ distance alone.

In addition to distance, we have employed a 1D deep ResNet of 19 convolutional layers to predict 3-state secondary structure and backbone torsion angles *ϕ* and *ψ* from position specific scoring matrix (PSSM) generated by HHblits^25^. To predict a distribution function for torsion angles, we train our ResNet model by maximizing the following probability function.

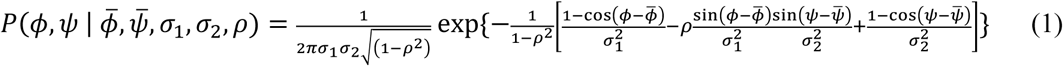

In Eqn.(1), 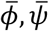 are the mean, *σ*_1_, *σ*_2_ are the variance and *ρ* is the correlation. That is, our deep ResNet outputs the mean and variance of the torsion angles at each residue.

### Multiple sequence alignment (MSA) and input features

We generated four different MSAs by running HHblits^25^ with 3 iterations and E-value set to 0.001 and 1 and running Jackhmmer^26^ with E-value set to 0.001 and 0.00001, respectively. HHblits searches through the uniclust30 library released in 2017 and Jackhmmer searches through the uniprot protein sequence database released in 2018. Metagenomics sequence database was not used for MSA generation. From each individual MSA, we derive protein sequential and pairwise features. Sequential features include sequence profile, secondary structure and solvent accessibility predicted by RaptorX-Property^27^. Pairwise features include standard and APC-corrected mutual information, pairwise contact potential and direct co-evolution strength calculated by CCMpred^28^. In summary, for each target we generated 4 sets of input features and accordingly 4 different distance predictions, which are then averaged to obtain the final prediction.

### Training data

We constructed our training and validation sets from PDB25 created early in 2018, which in total has 11410 proteins. No two proteins in this set share more than 25% sequence identity. We randomly selected about 900 proteins to form the validation set and used the remaining to form the training set. We have trained three models for each atom pair, which are then combined to form the final model.

### RaptorX-TBM

RaptorX-TBM used our new threading program DeepThreader^22^ to build sequence-template alignments and identify templates and then employed Rosetta-CM^29^ to build 3D models from alignment. DeepThreader greatly outperforms previous threading methods by integrating CNFpred^30^ with protein-specific distance potential predicted by our deep ResNet model. CNFpred is our old in-house threading program that aligns sequence to templates by integrating sequence profile, predicted secondary structure and solvent accessibility via Conditional Neural Fields^31^, which is a combination of shallow convolutional neural network and linear-graph-based Conditional Random Fields. CNFpred is on average more sensitive than HHpred^32^, but much worse than DeepThreader. RaptorX-TBM and CNFpred used PDB90 as the template database while our deep ResNet models for distance and angle prediction were trained by PDB25.

### RaptorX-Contact

In CASP13 we registered RaptorX-Contact for both contact prediction and distance-based template-free modeling. RaptorX-Contact converts predicted distance distribution, secondary structure and backbone torsion angles into CNS restraints and builds 3D models by running CNS^33^, a software program for experimental protein structure determination. Given a matrix corresponding to the distance probability distribution for each atom pair, we pick 7L (L is sequence length) pairs with the highest predicted likelihood (probability) having distance <15Å and assume that their distance is <15Å. From the predicted distance distribution of one atom pair, we estimate the mean distance *m* and standard deviation s, and then use *m-s* and *m+s* as its distance lower and upper bounds. We used the same method as CONFOLD^34^ to derive hydrogen-bond restraints from predicted alpha helices. Different from CONFOLD that derives torsion angles from predicted secondary structure, we use the mean degree and variance predicted by our 1D deep ResNet as torsion angle restraints.

For each protein, we run CNS to generate 200 possible 3D models and then choose 5 with the least violation of distance restraints as the final models. CNS uses distance geometry to build initial 3D models from distance restraints and then employs simulated annealing to refine bonds and angles so that the resultant models are protein-like. CNS can generate a 3D model very quickly. We generated multiple models for a protein since the CNS solution may not be globally optimal.

### RaptorX-DeepModeller

RaptorX-DeepModeller is also a distance-based folding server, differing from RaptorX-Contact in that the ResNet model used by RaptorX-DeepModeller has a few additional input features extracted from sequence-template alignment generated by RaptroX-TBM. The additional features include sequence-template similarity score (e.g., amino acid similarity, sequence profile similarity and secondary structure similarity) and an initial distance matrix extracted from the weakly similar template according to the alignment. Supposing two target residues i and j are aligned to two template residues k and l, we assign the distance between k and l as the initial distance of i and j. When one target residue is not aligned, the corresponding row and column in the initial distance matrix is empty.

RaptorX-DeepModeller is developed to study if we can improve protein structure modeling by integrating alignment, template and co-evolutionary information through deep learning. Due to the limit of computing resources, RaptorX-DeepModeller used in CASP13 was trained by the training set described in^6^, which is smaller than what was used for RaptorX-Contact in CASP13.

## Results

### Contact and distance prediction accuracy in CASP13

RaptorX-Contact was officially ranked first among 46 human and server contact predictors, in terms of a combination of several metrics. When top L/5, L/2 and L long-range predicted contacts are evaluated, on the FM targets RaptorX-Contact has precision 70.054%, 57.787% and 44.731%, respectively, and F1 values 0.233, 0.362 and 0.411, respectively. The other top 4 groups (which also used deep ResNet) have top L/5 long-range precision 65.678%, 64.031%, 60.798% and 60.595% and F1 values 0.213, 0.208, 0.192 and 0.191, respectively. As a control, MetaPSICOV^35^ (the CASP11 winner built upon a shallow neural network) ran by the CASP13 organizers has top L/5 long-range precision=25.16% and F1=0.078, respectively, and GaussDCA^36^ (the only DCA method blindly tested in CASP13) has precision=21.757% and F1=0.067, respectively. Fig. 2A and 2B show that the F1 value and precision of our deep ResNet method is not strongly correlated with MSA depth (trendline R^2^<0.38), which is different from the pure DCA method that heavily depends on MSA depth. Note that here we use the MSA depth downloaded from the CASP13 contact assessment web page.

**Figure 2.**
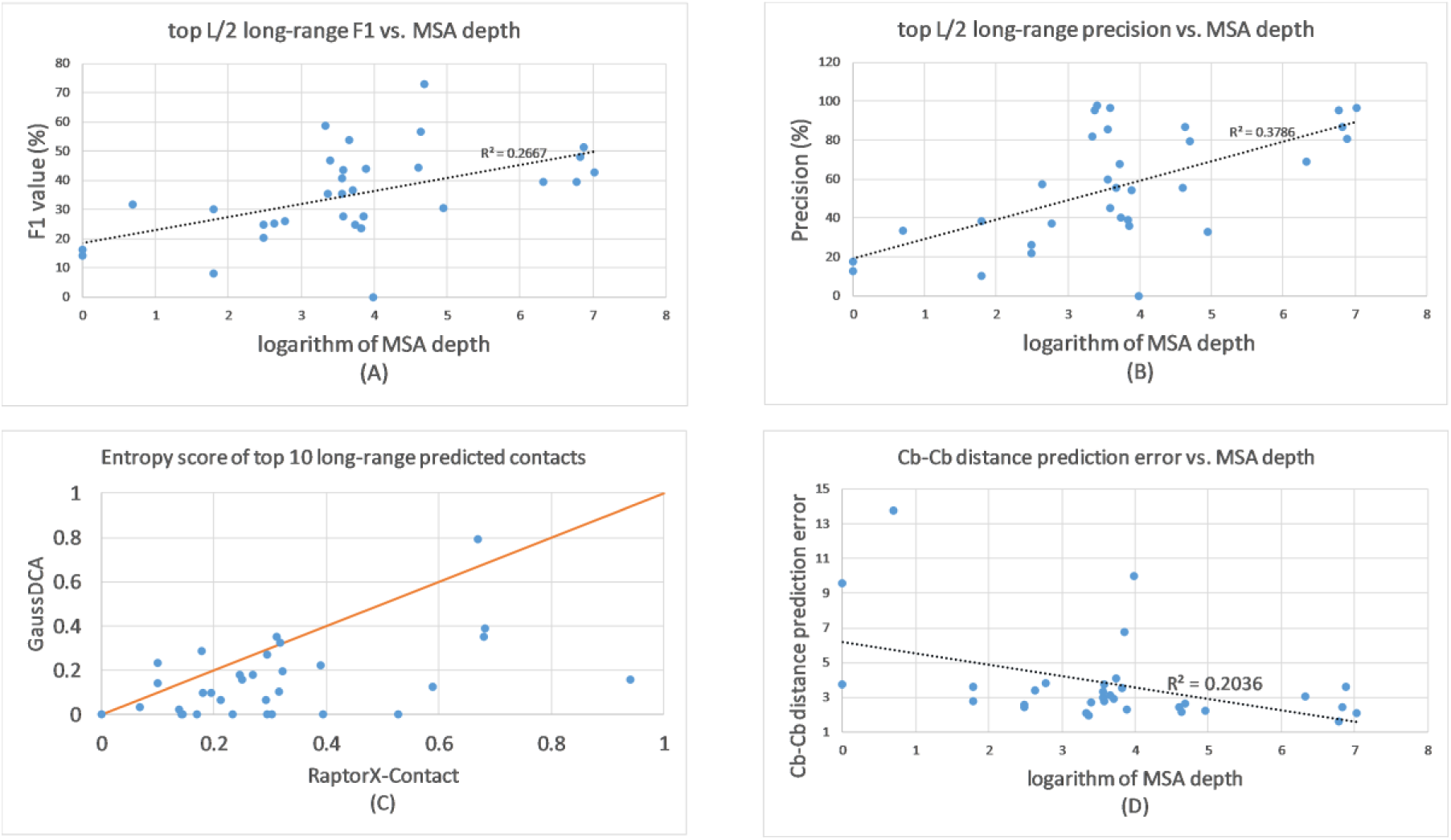
Contact and distance prediction analysis. (A) relationship between contact prediction F1 value and MSA depth; (B) relationship between contact prediction precision and MSA depth; (C) Entropy score of top 10 long-range contacts predicted by our method vs. GaussDCA; (D) relationship between Cb-Cb distance prediction error and MSA depth.

AlphaFold did not submit contact prediction, according to its presentation at the 7^th^ CAPRI meeting, it has a similar F1 value as RaptorX-Contact. On the top L/5, L/2 and L long-range contacts predicted for the FM targets, AlphaFold has F1 values 0.227, 0.369 and 0.419, respectively. That is, there is almost no performance difference between the deep network architectures used by AlphaFold and RaptorX-Contact.

In addition to precision and F1, entropy score is introduced by the contact prediction assessors in CASP12 to measure the spread-out of predicted contacts. A contact prediction with a large F1 value may not result in good 3D structure modeling if the predicted contacts are mainly located in a small contact submatrix. As reported by CASP13, when top 10, L/5 and L/2 long-range contacts are considered, RaptorX-Contact has entropy score 0.311, 0.643 and 1.255, respectively, much larger than GaussDCA^36^, which has entropy score 0.151, 0.332, and 0.553, respectively. Even when only top 10 contacts are considered, on most targets RaptorX-Contact has better entropy score than GaussDCA (Fig. 2C). This result indicates that contacts predicted by deep ResNet may contain more information content for 3D structure modeling than contacts predicted by DCA.

Our predicted C_β_-C_β_ distance on the FM targets has average error 3.76Å, precision 0.678, recall 0.540 and F1 0.588. While evaluating distance prediction, we consider only those atom pairs with sequence separation at least 12 and predicted distance <15Å. Recall is calculated as the ratio of atom pairs with native distance <15Å that are predicted to have distance <15Å. Precision is calculated as the ratio of atom pairs with predicted distance <15Å that have native distance <15Å. Our C_β_-C_β_ distance prediction error for most targets is less than 4Å (Fig. 2D) and it is not strongly correlated with MSA depth (coefficient=-0.45, trendline R^2^=0.2036). RaptorX-Contact predicted distance well for quite a few targets such as T0969-D1 (MSA depth>1000) and T0957s2-D1 (MSA depth 28), but did badly on T0953s1 and T0989-D1, both having MSA depth ∼50 (SI Appendix, Fig. S1). RaptorX-Contact failed on T0953s1 because it has only 34 long-range residue pairs with native distance <15Å, which is much smaller than typical. While estimating distance bounds from predicted distance distribution, we assumed each target had about 7L long-range pairs with distance<15Å, which resulted in a big prediction error. T0989 is a 2-domain target. Its 1^st^ domain has much better co-evolution signal than the 2^nd^ one. We did not split T0989 into 2 domains, which resulted in many more C_β_-C_β_ pairs in D1 being assumed to have distance<15Å and thus, led to a big prediction error. When T0989-D1 is predicted independently, its C_β_-C_β_ distance error is only 4.89Å.

### Distance-based tertiary structure modeling accuracy in CASP13

When all the CASP13 targets are considered, RaptorX-DeepModeller has the best 3D modeling performance among the three RaptorX servers. When only FM targets are considered, the average quality (TMscore) of the first models predicted by RatporX-DeepModeller, RaptorX-Contact and RaptorX-TBM are 0.471, 0.474 and 0.402, respectively. When all 5 models are considered for each target, RatporX-DeepModeller, RaptorX-Contact and RaptorX-TBM predicted correct folds (TMscore>0.5) for 17, 17, and 9 targets, respectively. On FM targets RaptorX-DeepModeller has slightly worse accuracy than RaptorX-Contact because the former was trained by a smaller training set than the latter. When combined together, RaptorX-DeepModeller and RaptorX-Contact predicted correct folds for 18 of the 32 FM targets.

Fig. 3 compares the performance of the three RaptorX servers and an old in-house threading program CNFpred on the FM targets. RaptorX-DeepModeller modeling accuracy is highly correlated with RaptorX-Contact (Fig. 3A) and RaptorX-TBM (Fig. 3B) since they mainly depend on pairwise distance information predicted by deep ResNet. Although both used the same template database, RaptorX-TBM performs much better than CNFpred (Fig. 3D) since RaptorX-TBM can recognize structurally similar templates even if they are not evolutionarily related to a target. This indicates the importance of pairwise distance in protein threading with remotely related templates. We did not have a contact-based threading program, but Zhang’s CEthreader is such a program and was tested in CASP13. RaptorX-TBM outperformed CEthreader by about 14% in terms of the TMscore of the first models on hard targets. This may suggest that predicted distance is much more informative than contacts for template-based modeling.

**Figure 3.**
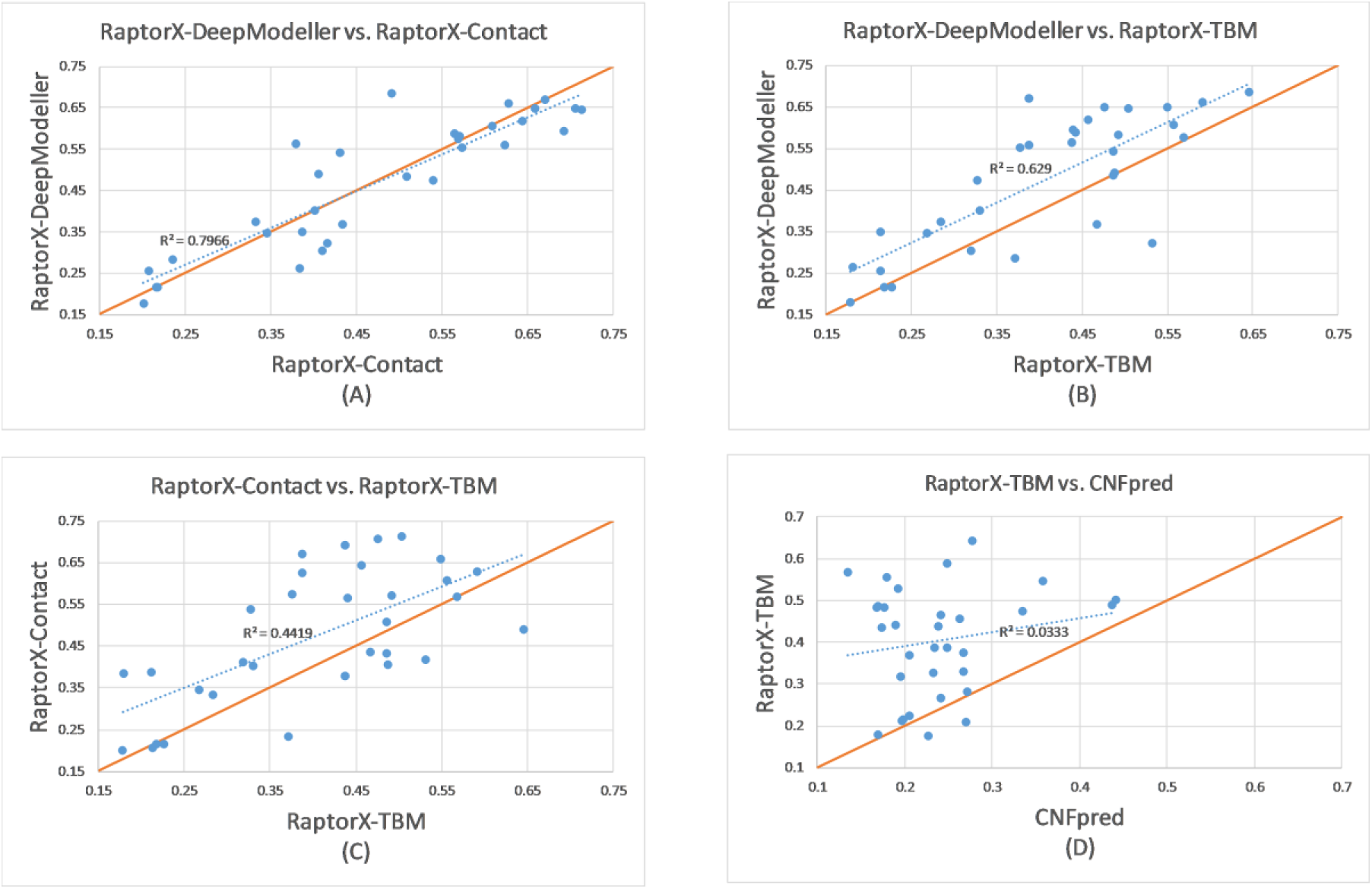
Comparison of 3D structure modeling accuracy of three RaptorX servers (RaptorX-DeepModeller, RaptorX-Contact and RaptorX-TBM) and an old in-housing threading program CNFpred.

RaptorX-DeepModeller and RaptorX-Contact clearly outperform RaptorX-TBM (Fig. 3C), which implies that for FM targets template-based modeling is insufficient even if predicted distance information is used to align sequence to templates and to select templates. For easier targets, RaptorX-DeepModeller has a larger advantage over both RaptorX-Contact and RaptorX-TBM. For example, among the 13 FM/TBM targets, RaptorX-DeepModeller predicted better models for 8 of them than RaptorX-Contact and RaptorX-TBM combined (Table 1). This may suggest that it is useful to integrate template and co-evolutionary information for structure modeling.

**Table 1.** The accuracy (TMscore) of the first models generated by RaptorX-DeepModeller, RaptorX-Contact and RaptorX-TBM on the 13 FM/TBM models.

| Target | Length | DeepModeller | RX-Contact | RX-TBM |
| --- | --- | --- | --- | --- |
| T0949-D1 | 129 | 0.687 | 0.631 | 0.685 |
| T0953s2-D1 | 44 | 0.180 | 0.166 | 0.155 |
| T0955-D1 | 41 | 0.589 | 0.344 | 0.688 |
| T0958-D1 | 77 | 0.657 | 0.554 | 0.609 |
| T0970-D1 | 85 | 0.535 | 0.547 | 0.458 |
| T0978-D1 | 413 | 0.701 | 0.583 | 0.680 |
| T0981-D3 | 203 | 0.650 | 0.545 | 0.637 |
| T0986s1-D1 | 92 | 0.689 | 0.655 | 0.654 |
| T0992-D1 | 107 | 0.745 | 0.745 | 0.712 |
| T0997-D1 | 185 | 0.774 | 0.757 | 0.577 |
| T1005-D1 | 326 | 0.705 | 0.641 | 0.677 |
| T1008-D1 | 77 | 0.280 | 0.276 | 0.278 |
| T1019s1-D1 | 58 | 0.376 | 0.441 | 0.327 |

### Progress in contact prediction and 3D structure modeling

We did not keep our old ResNet models developed in 2016, so cannot measure their performance on the CASP13 FM targets. Here we measure our progress using the 37 CASP12 FM targets. We compare contact prediction and folding accuracy of the two old versions of our ResNet model (denoted as CASP12-submit and CASP12-postdict) and the version trained right before CASP13 (denoted as CASP13-submit). CASP12-submit represents our model used during CASP12 in 2016. As explained before^9^, our ResNet method was under development during CASP12 and CASP12-submit was updated from time to time, so CASP12-submit is not a single complete version of our ResNet model. CASP12-postdict was trained right after CASP12, representing a full implementation of our deep ResNet method described in^6^. CASP13-submit was trained right before CASP13, improving over CASP12-postdict in the following aspects: 1) dilated convolution is used in CASP13-submit; and 2) 25 discrete distance bins are used in CASP13-submit as labels while 3 distance bins (0-8Å, 8-15Å and >15Å) are used in CASP12-postdict. To compare these three versions of our method, we use the same input features for the 37 CASP12 FM targets, which were generated in CASP12. Note that here for fair comparison, both CASP12-postdict and CASP13-submit were trained by the same set of ∼10,000 training proteins. Nevertheless, while used in CASP13, CASP13-submit was trained by a slightly larger set of 11410 training proteins.

CASP12-postdict yields much better contact prediction and structure modeling than CASP12-submit because the former is a complete implementation of our ResNet method while the latter is not (Table 2). The results of both CASP12-postdict and CASP12-submit are taken from Tables 1 and 3 of our previous paper^9^. For contact prediction, on average CASP13-submit outperforms CASP12-postdict by about 6-7%. That is, compared to our CASP12 submission, we have greatly improved contact prediction, but compared to the ResNet model we have fully implemented in 2016, the improvement is only 6-7%. The major improvement of our CASP13 method over CASP12-postdict lies in 3D structure modeling. Table 2 shows that by using distance-based instead of contact-based folding, we may improve 3D structure model quality by 0.1 in terms of TMscore. For most targets, distance-based folding generated better 3D models than contact-based folding (Fig. 4).

**Table 2.** Progress in terms of long-range contact prediction precision and 3D structure modeling on the 37 CASP12 FM targets.

|  | Long-range contact precision (%) |  |  |  | Folding accuracy (TMscore) |  |
| --- | --- | --- | --- | --- | --- | --- |
|  | L | L/2 | L/5 | L/10 | Top 1 | Top 5 |
| CASP12-submit | 28.63 | 36.42 | 46.76 | 51.50 | 0.274 | 0.307 |
| CASP12-postdict | 40.18 | 50.20 | 58.87 | 63.93 | 0.354 | 0.397 |
| CASP13-submit | 43.10 | 56.90 | 66.90 | 73.80 | 0.466 | 0.476 |

**Figure 4.**
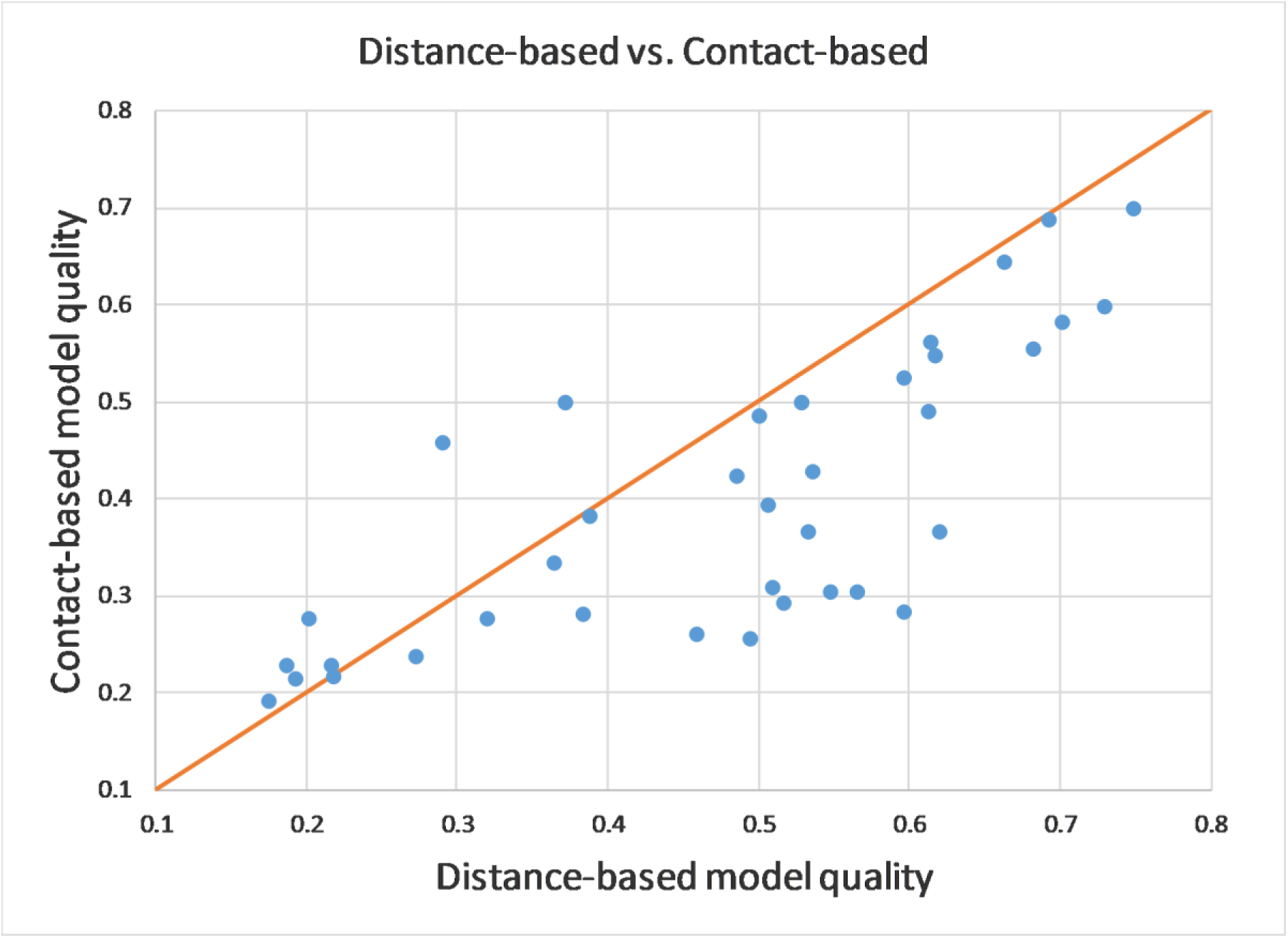
Distance-based folding accuracy vs. contact-based folding accuracy, measured on the 37 CASP12 FM targets.

### Case Study

In this section, we study the models predicted by our servers for the three largest CASP13 FM targets: T0950-D1, T0969-D1 and T1000-D2. They have 342, 354, and 368 residues with valid coordinates, respectively, and MSA depth 103, 1132 and 877, respectively. Our servers predicted correct folds for all three targets and the best models for T0950-D1 and T0969-D1 among all human and server groups. In contrast, AlphaFold predicted correct folds for two of them, although on average AlphaFold has better modeling accuracy on all the FM targets.

#### T0950-D1

This is a very hard target and the ratio between its MSA depth and sequence length is only ∼0.3. Among all the 3D models accepted by CASP13, only seven have a correct fold (i.e., TMscore>0.5), including 5 models generated by RaptorX-DeepModeller, one by Zhang’s QUARK^37^ and one by Cheng’s human group MULTICOM. The best model was generated by RaptorX-DeepModeller, which has TMscore=0.589 (Fig. 5). MULTICOM’s best model is a copy of RaptorX-DeepModeller’s first model with TMscore=0.564. Zhang’s and AlphaFold’s best models have TMscore 0.506 and 0.443, respectively. Note that although this is a server-only target, some human groups such as AlphaFold and MULTICOM still submitted their predictions. Structure alignment by DeepAlign shows that the most similar training protein in our training set has TMscore=0.542 with this target. The template-based models predicted by RaptorX-TBM and CNFpred have TMscore=0.437 and 0.173, respectively, much worse than RaptorX-DeepModeller models. This implies that RaptorX-DeepModeller predicted models for this target not by only copying from a single template, although RaptorX-DeepModeller used alignments generated by RaptorX-TBM as input. That is, RaptorX-DeepModeller is able to generate better models than both RaptorX-Contact and RaptorX-TBM by combining alignment, template and co-evolution information.

**Figure 5.**
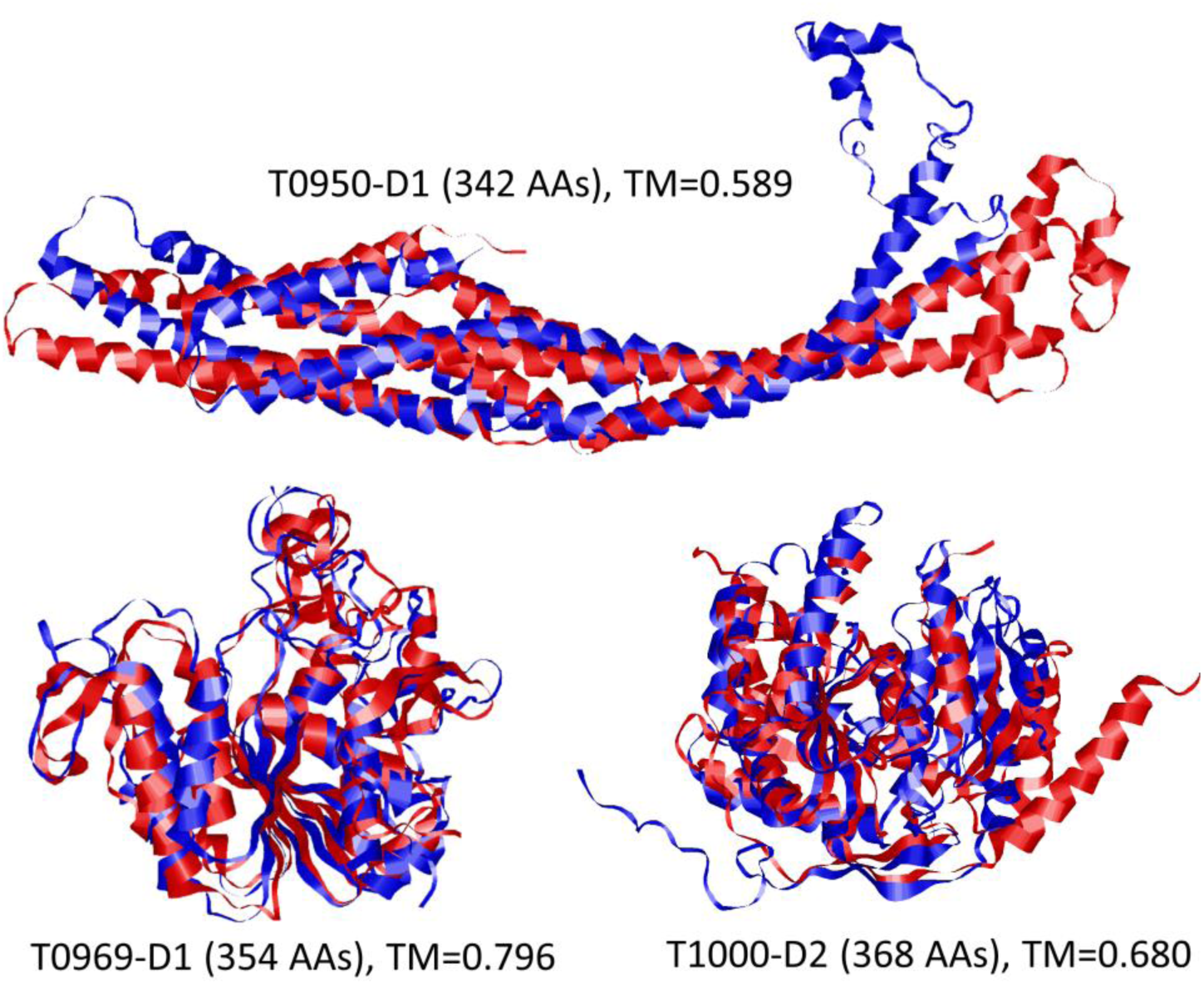
Superimposition between the best 3D models (blue) predicted by our servers and the native structure (red) for three largest CASP13 FM targets (T0950-D1, T0969-D1 and T1000-D2).

#### T0969-D1

This target has the largest MSA depth among all CASP13 FM targets. Quite a few groups predicted models with a correct fold. RaptorX-Contact predicted a model with TMscore=0.796 (Fig. 5), better than all the other server models and all but 4 human models (MESHI, MUFold, Seder3mm and Elofsson), which have similar quality as the RaptorX-Contact model simply because they are derived from this server model. For this target, AlphaFold’s and Zhang’s best models have TMscore=0.730 and 0.680, respectively. The most similar training protein in our training set has TMscore=0.452 with this target. The template-based models predicted by RaptorX-TBM and CNFpred have TMscore=0.549 and 0.357, respectively. This implies that our server model is not copied from individual proteins in our training set or template database.

#### T1000-D2

This is a very large protein domain with MSA depth being 877. Quite a few groups predicted models of a correct fold. In particular, AlphaFold and Zhang predicted very good models with TMscore=0.880 and 0.851, respectively. Our servers have also predicted a correct fold (Fig. 5), but with much lower model quality (TMscore=0.680). The most similar training protein in our training set has TMscore=0.347 with the target. The template-based models predicted by RaptorX-TBM and CNFpred have TMscore=0.387 and 0.233, respectively. That is, our server model is not copied from individual proteins in our training set or template database. For this target, our contact prediction is ranked very top, better than Zhang’s contact prediction regardless of ranking metrics, but our 3D modeling has much lower quality than Zhang’s. This may indicate that we did not do a good job in building 3D model from predicted distance distribution.

## Discussion and Conclusion

Our CASP13 result confirms that by using deep ResNet to predict inter-atom or inter-residue distance, we may fold proteins much more accurately than ever before on a Linux workstation of 20 CPUs within minutes to hours. In particular, our distance-based folding algorithm predicted the best models for two of the three largest FM targets with ∼350 residues. Our analysis shows that predicted distance is useful for both template-free and template-based modeling, and that protein modeling can be further improved by combining template and co-evolutionary information.

### What went right?

The CASP13 result is consistent with our previous findings: (1) protein distance matrix can be predicted very well by a deep and global ResNet; (2) predicted distance can greatly improve both template-based and template free modeling; (3) predicted distance is more informative than contacts for protein structure modeling; (4) protein structure modeling can be further improved by integrating template and co-evolutionary information. Our protocol for contact prediction works very well even if we did not use as many layers as AlphaFold or as many input features as other top groups. Our distance geometry method for building 3D models from predicted distance works fine on very large targets even without time-consuming conformation sampling.

### What went wrong?

Although our protocol for predicting contacts from predicted distance distribution worked very well, our protocol for building 3D models from predicted distance distribution was not optimal. Our contact prediction has similar F1 value as AlphaFold, but our 3D modeling protocol underperformed AlphaFold on quite a few FM targets even if on very large targets our method has favorable performance. This may imply that for many FM targets we did not do a good job in deriving distance bounds from predicted distance distribution. One possible issue is that while estimating mean distance and standard deviation from predicted distance distribution, we simply ignored the predicted probability of distance >16Å for those atom pairs which are likely to have native distance less than 15Å. The other possible issue is that we assumed that each target has at least 7L C_β_-C_β_ pairs with native distance less than 15Å. Such an assumption leads to a big distance prediction error for T0953s1, which has only a small number of C_β_-C_β_ pairs with native distance <15Å.

We did not handle some multi-domain targets very well, especially when domains have very different MSA depth. In this case some domains of a target may have much stronger co-evolution signal than the others. Since we did not split a multi-domain target into domains when none of them have reasonable templates, the atom pairs chosen by us to build 3D models may not spread uniformly across different domains or segments. As such, some domains (or segments) may be covered by very few selected atom pairs while others may be covered by many more selected atom pairs. Both scenarios may lead to bad 3D modeling.

Our current folding protocol does not use fragment assembly, sophisticated energy functions or time-consuming conformation sampling, which may prevent us from generating high-resolution good models. For example, some of our models for beta sheets do not form very good hydrogen-bonding since our folding method does not have any energy terms promoting the formation of beta sheet. Finally, we have not used the metagenomics sequence database for MSA generation, which may be helpful for few targets.

### Why did deep learning perform much better in CASP13 than before?

Deep learning such as Deep Belief Networks (DBN) has been attempted for protein contact prediction in 2012^38, 39^, but it drew little attention from the community. This is mainly because that the DBN method has almost the same performance as traditional machine learning methods such as Random Forests, as reported by the developers^38, 39^, who have tested this method in CASP10, CASP11 and CASP12. In CASP11 and CASP12, the DBN method even underperformed MetaPSICOV, a traditional neural network method. The main reason why deep learning becomes effective for protein folding in the past couple of years is not the enlargement of protein sequence databases, but the introduction of a totally new formulation of contact prediction (i.e., simultaneous prediction of all contacts in a protein) and new network architecture (i.e., deep and global ResNet). In fact, the CASP13 FM targets have similar MSA depth as the CASP12 FM targets, but the contact prediction and folding accuracy on CASP13 FM targets is much higher due to community-wide adoption of deep and global ResNet for contact and distance prediction.

The deep ResNet method is not simply an enhancement of the DBN method. Their difference is analogous to that between DCA (direct coupling analysis) and mutual information. Both DCA and deep ResNet are global methods while mutual information and DBN are local methods. While predicting the label (i.e., contact or distance) of two residues, global methods look at the labels of all other residue pairs, but local methods do not. By applying a global convolutional operation to the whole contact/distance matrix, deep ResNet may learn protein structure patterns easily and yield much better contact/distance prediction. As pointed out by us before^9^, when one contact is predicted independent of the others, even if ResNet is applied, the resultant contact prediction accuracy is not very good. Further, it is much easier to build a very deep ResNet than a very deep DBN, which limits the performance of DBN.

Our ResNet method for contact prediction (i.e., RaptorX-Contact) was not fully implemented during CASP12, not to mention the folding protocol. Because of this, our CASP12 contact prediction accuracy is not much higher than the other groups, although RaptorX-Contact was still ranked first in CASP12. As shown in Table 2 and previous blind CAMEO test^40^, a full implementation of our ResNet method right after CASP12 has much better contact prediction and folding accuracy.

### The major difference among top groups in CASP13

Many CASP13 groups have incorporated deep convolutional neural network into their structure prediction pipelines. A nature question to ask is what are their major difference? Here we analyze 4 representative groups: AlphaFold, Zhang, RaptorX and Baker^41^. We do not consider some top human groups such as Cheng’s MULTICOM since they heavily relied on consensus analysis of server models, and it is unclear if there are any major methods underlying their results other than consensus analysis.

AlphaFold used deep ResNet to predict inter-residue distance distribution, converted this distribution to protein-specific distance potential and then minimized it by conformation sampling and gradient-based methods to build 3D models. Zhang used ResNet to predict inter-residue contacts and then used them to guide folding simulation. RaptorX used ResNet to predict inter-atom distance distribution, converted it to mean distance and deviation and then used this as distance bounds to build 3D models by CNS. Baker used DCA to predict contacts for some targets and then conducted contact-assisted folding simulation. On average Baker’s Rosetta underperformed the other three groups because of lack of the deep learning module, although Rosetta can do extensive conformation sampling. AlphaFold and RaptorX have a similar contact prediction performance, which is slightly better than Zhang. In terms of 3D modeling, RaptorX has similar performance as Zhang’s servers, although their methods seem to be different. For 3D modeling, RaptorX’s strength lies in distance prediction, which provides more informative than contact prediction, but Zhang made it up by incorporating predicted contacts into a well-developed folding engine. Compared to RaptorX, AlphaFold has a much better folding protocol, i.e., building 3D models by conformation sampling and gradient-based optimization methods. Compared to Zhang, AlphaFold has more informative distance prediction. In summary, AlphaFold did well in both the deep learning and model building steps while the other three groups did well in only one of them, which is why AlphaFold stands out in modeling FM targets.

## Acknowledgement

This work is supported by National Institutes of Health grant R01GM089753 to JX and National Science Foundation grant DBI-1564955 to JX. The funders had no role in study design, data collection and analysis, decision to publish, or preparation of the manuscript. The authors thank CASP13 organizers and assessors and all contributors of the experimental data.

## Author Contributions

JX conceived, designed and implemented the major algorithms and wrote the paper. SW built the initial pipeline for MSA generation and template-based modeling and helped draw Figure 1.

## Supporting Information

**Figure S1.**
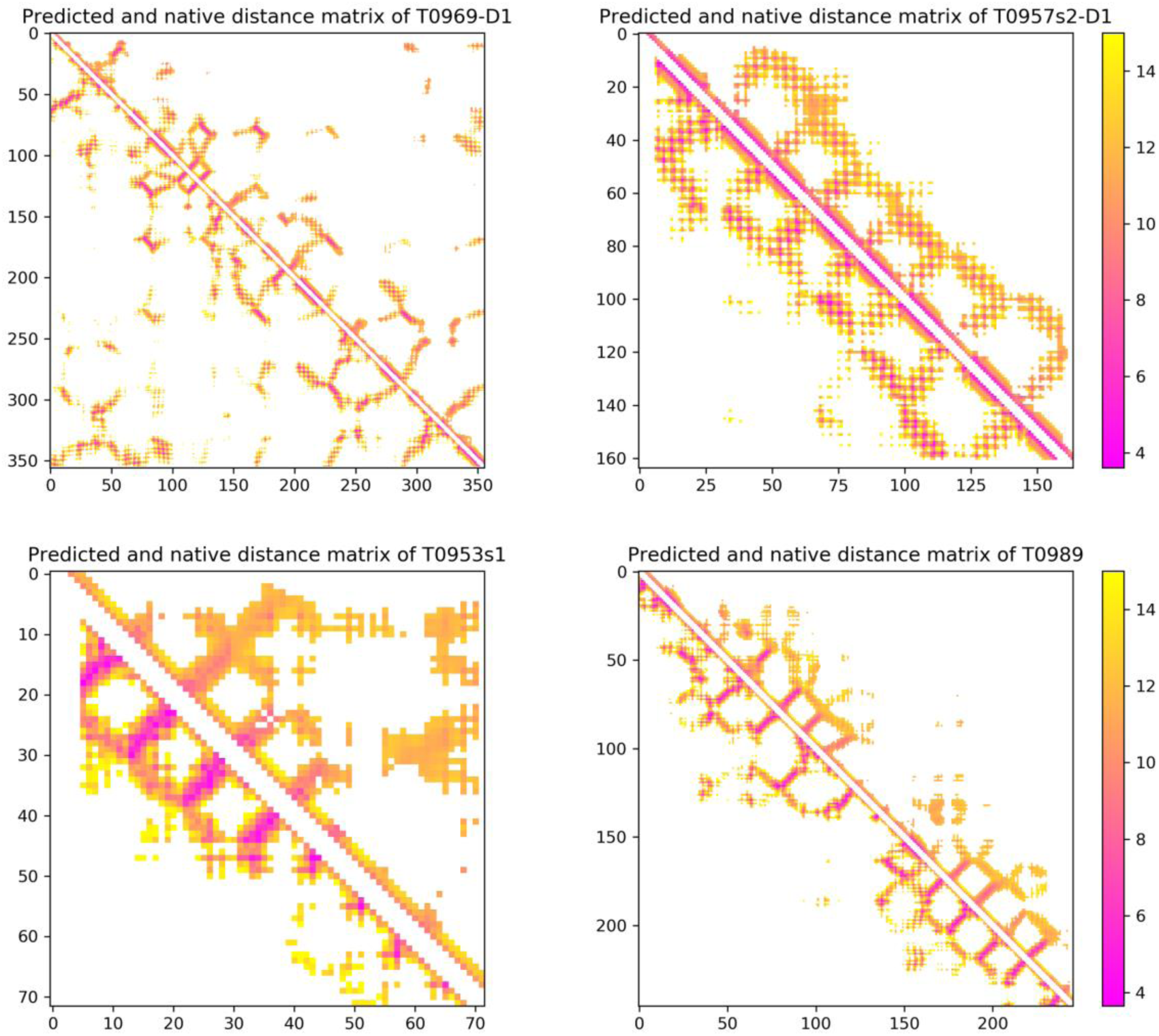
Native (lower triangle) and predicted (upper triangle) distance matrices of 4 CASP13 hard targets: T0969-D1, T0957s2-D1, T0953s1-D1 and T0989-D1.

